# ClaroAI-Bench: Evaluating Agentic Scientific Reproducibility on Real Biomedical Papers

**DOI:** 10.64898/2026.05.08.723611

**Authors:** Kyle O’Connell

## Abstract

We introduce **ClaroAI-Bench**, an evaluation suite for measuring AI agents’ ability to reproduce computational findings from published biomedical research. The benchmark comprises 35 real NIH-funded papers spanning five modalities (genomics, imaging, clinical/EHR, epidemiology, wet-lab) scored on a five-dimension rubric: data findability (D1), data accessibility (D2), code availability (D3), environment reconstructability (D4), and results reproducibility (D5). Each task requires an agent to locate data, obtain code, reconstruct the compute environment, execute the analysis, and verify results against published claims—mirroring the full scientific reproduction pipeline. In a three-condition ablation, an audit-only baseline (D1–D4 metadata scoring) and a bash-only agent (API + bash tool) both achieve 0% D5 reproduction, while a full-capability agent (Claude Code, all tools) reproduces 20 of 33 computational papers (60.6%; 95% CI [42.4, 75.8]). D1–D4 metadata scores strongly predict D5 outcomes (Spearman *r*=0.68, *p*<0.0001), and papers with accessible data and code achieve 2.9× higher D5 scores than restricted papers (*p*=0.0013). Multi-model scoring with three frontier models (Claude Opus 4.6, GPT-5.4, Gemini 2.5 Pro) yields inter-model agreement of *r*=0.85–0.97 on D3 but only *r*=0.51–0.81 on D4, identifying environment reconstruction as the dimension with highest evaluator disagreement. ClaroAI-Bench fills a gap between code-generation benchmarks (SWE-bench) and end-to-end scientific AI evaluations by testing long-horizon, real-world reproduction tasks with brittle environments, missing meta-data, and access constraints. The benchmark, scoring rubric, agent logs, and pip-installable auditor are archived at https://doi.org/10.5281/zenodo.20071236 and on HuggingFace Datasets at https://huggingface.co/datasets/kyleaoconnell22/claroai-bench.

## 1 Introduction

AI agents are increasingly applied to scientific tasks: code generation (Jimenez et al., 2024), literature review (Lu et al., 2024), and experiment design (Boiko et al., 2023). Yet no benchmark directly evaluates whether an agent can *reproduce* a published scientific result end-to-end—the gold standard of the scientific method. Existing agent benchmarks test code repair (SWE-bench (Jimenez et al., 2024)), AI research replication (PaperBench (Starace et al., 2025)), social-science reproducibility assessment (REPRO-Bench (Hu et al., 2025)), or end-to-end physics paper reproduction (PRBench (Qiu et al., 2026)), but none require the agent to navigate the full biomedical reproduction pipeline: locating data in heterogeneous repositories, obtaining code that may be broken or absent, reconstructing compute environments from incomplete specifications, executing analyses that span languages and frameworks, and quantitatively verifying results against published claims.

ClaroAI-Bench fills this gap with 35 real biomedical reproduction tasks drawn from NIH-funded research published in 2025–2026. Each task is defined by a published paper and its associated artifacts (data deposits, code repositories, methods descriptions) rather than by synthetic test cases, ensuring that tasks reflect the actual difficulty distribution of scientific reproducibility. The benchmark makes three contributions:

1. **A graded, multi-dimensional evaluation rubric**. Each paper is scored on five dimensions (D1– D5, 0–2 each), providing fine-grained signal about *where* reproduction fails—not just whether it succeeds.
2. **An ablation separating tool access from capability**. We compare audit-only (no execution), bash-only (single-tool agent), and full-capability (all tools) conditions, showing that tool access is necessary but not sufficient for reproduction.
3. **Multi-model scoring for rubric validation**. Three frontier models independently score the same papers, yielding inter-model reliability statistics that identify which rubric dimensions are operationally unambiguous and which require calibration.

Beyond benchmarking AI agents, ClaroAI-Bench provides the first large-scale empirical measurement of computational reproducibility under the NIH Data Management and Sharing (DMS) Policy, revealing that data *discoverability* has improved dramatically (D1 mean = 1.69/2) while end-to-end *reproducibility* remains challenging (D5 mean = 1.03/2, failure rate 39.4%).

## 2 Related Work

### Agent benchmarks

SWE-bench (Jimenez et al., 2024) tests agents on GitHub issue resolution (2,294 Python tasks); WebArena (Zhou et al., 2024) evaluates web browsing; MLE-bench (Chan et al., 2024) tests Kaggle competition performance. These benchmarks focus on well-defined, single-domain tasks. Scientific reproduction is harder: it is multi-step, multi-domain, and often under-specified. PaperBench (Starace et al., 2025) evaluates AI agents on replicating 20 ICML papers from scratch; REPRO-Bench (Hu et al., 2025) evaluates reproducibility assessment of social-science papers given reproduction packages; PRBench (Qiu et al., 2026) studies end-to-end paper reproduction on 30 physics tasks. ClaroAI-Bench differs by focusing on real biomedical papers and by grading failures at each stage of the reproduction pipeline rather than only final task completion.

### Reproducibility studies

The Reproducibility Project: Cancer Biology (Errington et al., 2021) replicated 50 experiments across 5 years at $2M+ cost. Surveys of computational results also document substantial manual burden from data acquisition, environment reconstruction, and code debugging (Stodden et al., 2018). Our work combines benchmark-style evaluation with real reproduction, bridging the AI evaluation and metascience communities.

### NIH DMS Policy evaluation

The NIH DMS Policy (NOT-OD-21-013, effective January 2023) requires data sharing for all NIH-funded research. To our knowledge, no prior study has measured whether this policy translates into computational reproducibility. ClaroAI-Bench provides the first automated assessment.

## 3 Benchmark Design

### 3.1 Task Definition

Each ClaroAI-Bench task is defined by a published biomedical paper and consists of five sequential stages:

1. **Extraction:** Parse the paper to identify data repositories, code links, key quantitative results, and statistical methods.
2. **Accessibility:** Download all referenced datasets and clone all code repositories.
3. **Environment:** Reconstruct the compute environment from dependency specifications or infer dependencies from code.
4. **Execution:** Run the analysis pipeline or, when code is absent, reconstruct it from the methods section.
5. **Comparison:** Compare regenerated results to published values using tolerance-based matching.

This five-stage pipeline mirrors how a human scientist would attempt reproduction, and each stage maps to a scoring dimension (D1–D5).

### 3.2 Scoring Rubric (D1–D5)

Each dimension is scored on a 0–2 scale:

- **D1: Data Findable** — 0: no data statement; 1: broken links or “upon request”; 2: valid repository with accession numbers.
- **D2: Data Accessible** — 0: download fails or gated; 1: partial access or registration; 2: fully downloadable.
- **D3: Code Available** — 0: no code; 1: partial code or detailed methods; 2: complete code with README.
- **D4: Environment Reconstructable** — 0: no dependency spec; 1: fixable build; 2: clean build from lockfile/Docker.
- **D5: Results Match** — 0: pipeline fails; 1: qualitative match; 2: quantitative match within 5%.

D4 and D5 are scored N/A for wet-lab papers (2 of 35), yielding a maximum of 10 points for computational papers and 6 for wet-lab. The rubric is defined in YAML configuration files shipped with the claroai package.

### 3.3 Paper Selection

Papers were selected using stratified random sampling from PubMed, restricted to NIH-funded publications from 2025–2026 with data availability statements. The initial 20 papers were sampled across five modalities (genomics, imaging, clinical/EHR, epidemiology, wet-lab) and two funding types: *intramural* (research conducted by scientists employed directly at NIH facilities under the NIH Intramural Research Program) and *extramural* (research conducted at universities or other institutions and funded by NIH grants). An additional 15 expansion papers were selected to enrich the “Public-Core” subset—papers with confirmed open code and data—enabling statistical comparison between open and restricted papers. Table 1 summarizes the benchmark composition.

**Table 1:** ClaroAI-Bench composition (35 papers by modality and NIH funding type). Of the 35 papers, 33 involve computational analyses and are scored on all five dimensions (D1–D5); the 2 wet-lab papers are scored on D1–D3 only.

| Modality | Intramural | Extramural | Total |
| --- | --- | --- | --- |
| Genomics/Omics | 3 | 7 | 10 |
| Imaging | 2 | 4 | 6 |
| Clinical/EHR | 2 | 3 | 5 |
| Epidemiology | 2 | 7 | 9 |
| Wet-lab | 1 | 1 | 2 |
| Mixed/Other | 0 | 3 | 3 |
| **Total** | **10** | **25** | **35** |

### 3.4 Difficulty Distribution

Tasks span a wide difficulty range. At the easy end, Paper 23 (TorchXRayVision) is a pip-installable library with auto-downloading model weights and machine-readable benchmarks (D1–D5 all = 2). At the hard end, Paper 03 requires three DUA-gated neuroimaging datasets (OASIS3, ADNI, AIBL) and proprietary software, making automated reproduction impossible (D1–D5 = [1,0,0,0,0]). This range ensures the benchmark discriminates between agents of varying capability.

## 4 Experiments

### 4.1 Three-Condition Ablation

We evaluate reproduction under three conditions on the 33 computational papers:

1. **Audit-only** (D1–D4 scoring): The agent scores metadata dimensions without executing any code. This measures what can be inferred from paper artifacts alone.
2. **Bash-agent**: An Anthropic API agent with access only to a bash tool. It can execute commands but cannot browse the web, read files intelligently, or spawn sub-agents.
3. **Full-agent**: Claude Code (Opus 4.6, 1M context) with all tools—bash, file I/O, web search, web fetch, code editing, and sub-agent spawning. This represents the current frontier of agentic capability.

### 4.2 Multi-Model D1–D4 Scoring

Three frontier models independently scored all 35 papers on D1–D4: Claude Opus 4.6 (Anthropic), GPT-5.4 (OpenAI), and Gemini 2.5 Pro (Google). Each model received an identical structured prompt containing the paper’s metadata, extracted repository URLs, HTTP accessibility results, and the D1–D5 rubric with level descriptors. No model was given tool access during scoring; scores were based solely on the structured audit metadata.

### 4.3 Implementation and Compute

The full pipeline is implemented in the `claroai` Python package (`pip install -e`.; 3,446 lines) with 31,430 lines of per-paper reproduction scripts across four languages (Python, R, MATLAB, SAS). Each agent run writes structured JSON to `papers/paper_XX/scores.json`. Compute-intensive D5 runs (WGS variant calling, large-scale data downloads) were executed on Google Cloud Platform VMs totaling 402 vCPU-hours.

#### Data scale

The 35 benchmark papers reference datasets totaling 8.3 TB at full scale, of which 1.7 TB is openly accessible without data use agreements. The largest datasets include 5,909 *M. tuberculosis* genomes from ENA (~850 GB FASTQ), TCGA/CPTAC whole-slide images for pathology evaluation (~500 GB), and the Genecorpus-30M single-cell corpus (~200 GB). Our agents processed 28.4 GB directly; the remainder was either DUA-gated (11 papers) or subsampled for tractability (e.g., 62 of 5,909 TB genomes).

#### Cost

The total cost of running ClaroAI-Bench was $112: $94 in LLM API calls (D1–D5 scoring, D5 full-agent reproduction, multi-model scoring, and 18 variance runs) and $18 in GCP compute. Direct human– AI cost comparisons are imperfect, but this level of spend is modest relative to prior multi-year human-led replication efforts in cancer biology and to survey evidence that computational reproduction routinely requires substantial manual debugging and data acquisition effort (Errington et al., 2021; Stodden et al., 2018).

All agent prompts, reproduction logs, and GCP pipeline scripts are included in the release.

**Figure 1:**
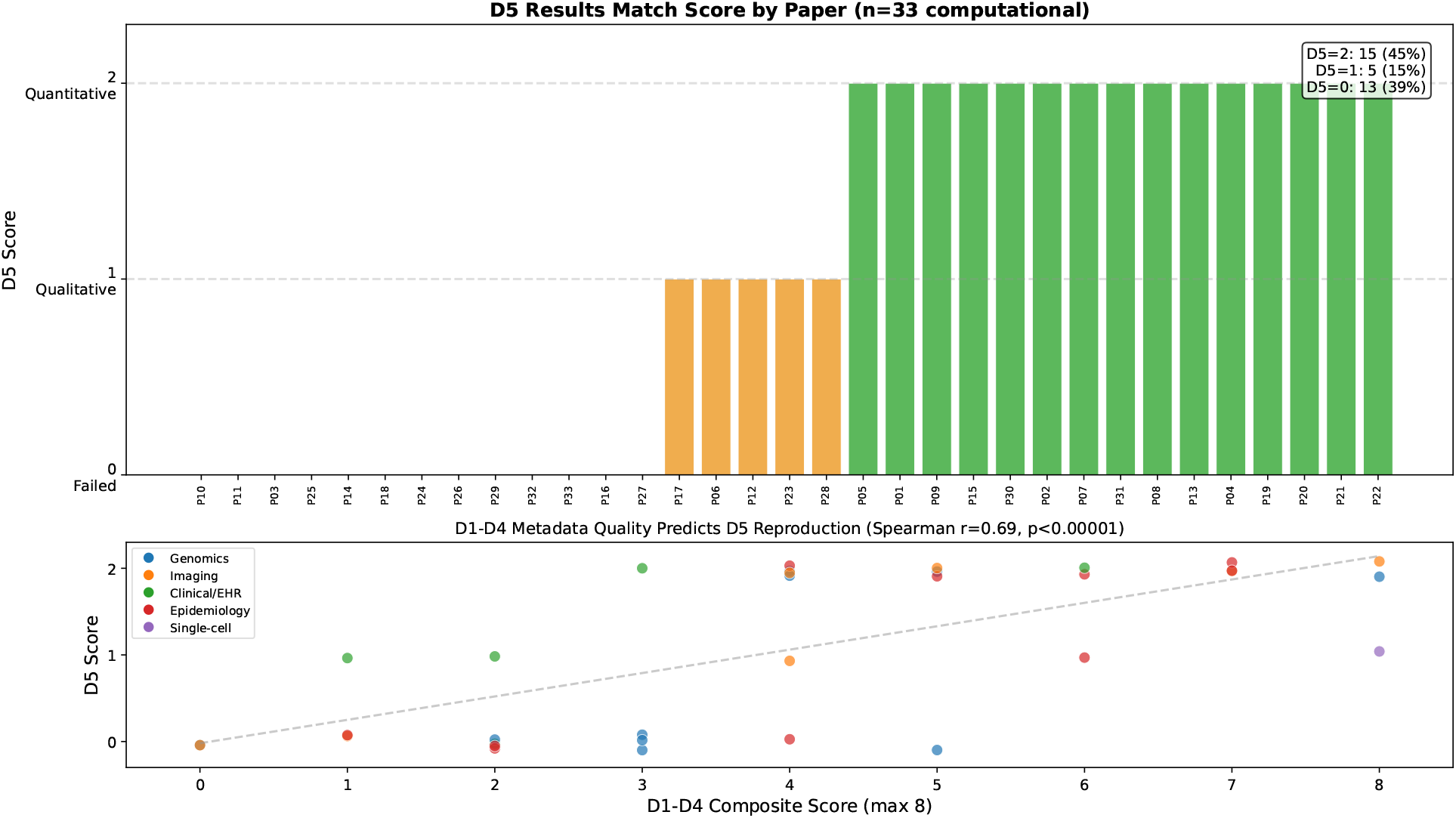
D5 reproduction outcomes across the 33 computational papers under the full-agent condition, ordered by score. Blue = D5=2 (quantitative match); orange = D5=1 (qualitative); red = D5=0 (failed or DUA-blocked). Papers with accessible data and code (Public-Core) cluster at the top.

## 5 Results

### 5.1 Ablation: Tool Access Is Necessary but Not Sufficient

The audit-only baseline achieves 0% D5 reproduction by construction—it cannot execute code. The bash-agent also achieves 0%, demonstrating that raw command-line access is insufficient without the ability to read documentation, search the web for download instructions, debug error messages, and adapt strategies. The full-capability agent reproduces 20 of 33 papers (60.6%; bootstrap 95% CI [42.4%, 75.8%]), with a D5 mean of 1.03/2.

**Table 2:** D5 reproduction rates by condition (33 computational papers).

| Condition | D5 > 0 | Rate | D5 Mean |
| --- | --- | --- | --- |
| Audit-only (D1–D4) | 0/33 | 0.0% | 0.00 |
| Bash-agent (API + bash) | 0/33 | 0.0% | 0.00 |
| Full-agent (Claude Code) | 20/33 | 60.6% | 1.03 |

### 5.2 D1–D4 Predict D5

Upstream metadata scores (D1–D4 composite) strongly predict D5 outcomes. Spearman correlation: *r* = 0.678, *p* = 1.4 × 10^−5^ (Figure 2). Papers in the “Public-Core” subset (D2 ≥ 1 and D3 ≥ 1, *n* = 15) achieve D5 mean = 1.60, with 93% (14/15) achieving D5 > 0. Restricted papers (*n* = 18) achieve D5 mean = 0.56 (Mann–Whitney *p* = 0.0013). This 2.9× gap confirms that reproducibility is primarily a function of *what artifacts are shared*, not agent capability.

**Figure 2:**
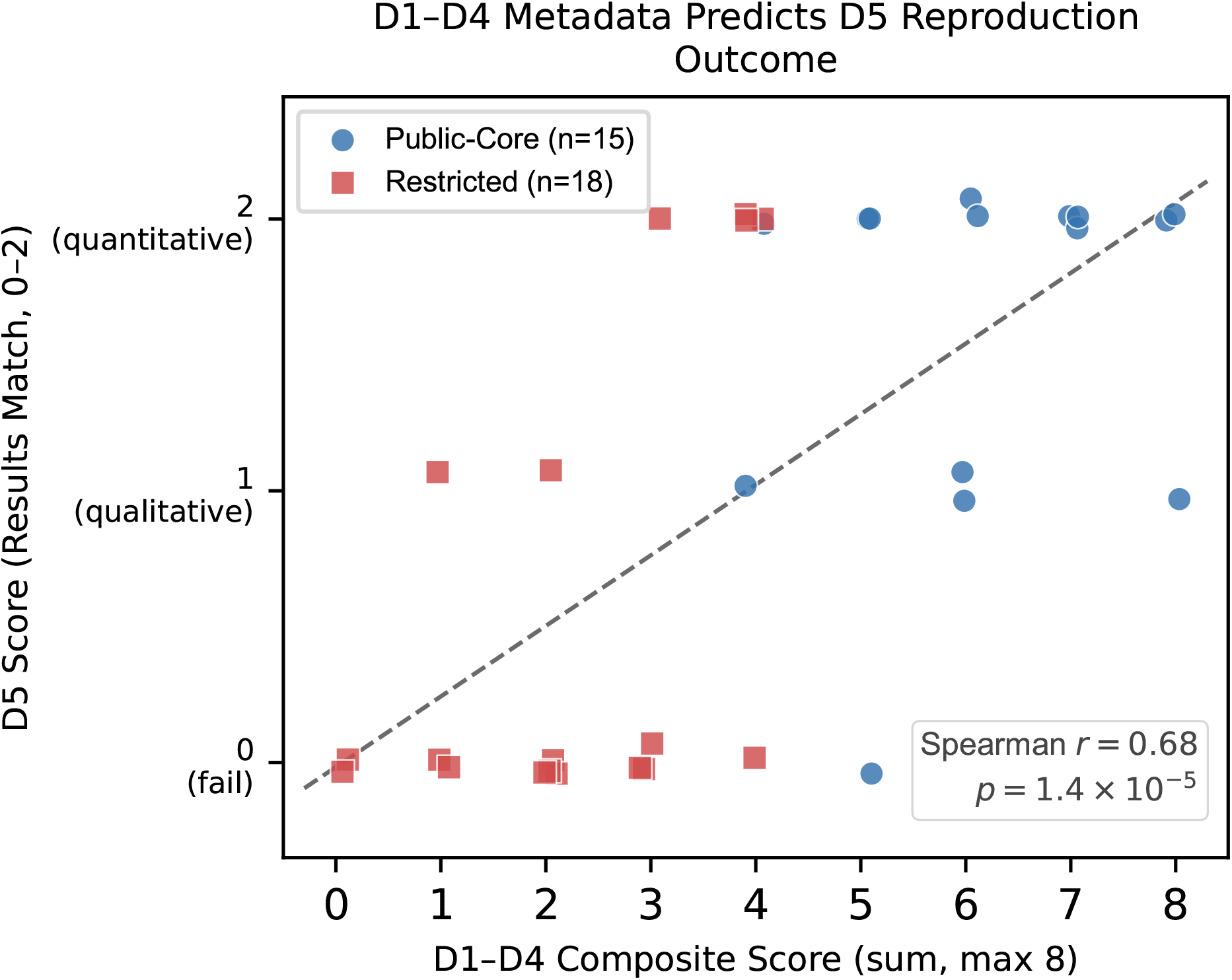
D1–D4 composite score predicts D5 reproduction outcome (Spearman *r*=0.678, *p*=1.4 × 10^−5^, *n*=33). Blue circles: Public-Core papers (D2 ≥ 1 and D3 ≥ 1); red squares: restricted papers. Dashed line: OLS fit.

### 5.3 Dimension-Level Analysis

The “staircase” pattern (Figure 3) shows data discoverability at near-ceiling (D1 failure rate 8.6%), dropping sharply through accessibility (42.9%), code availability (31.4%), environment reconstruction (57.6%), and results matching (39.4%). This cascade means that improving any single upstream dimension benefits all downstream dimensions.

**Table 3:** Per-dimension means and failure rates across 35 papers.

| Dimension | Mean | Max | Failure (score=0) | $n$ |
| --- | --- | --- | --- | --- |
| D1: Data Findable | 1.69 | 2 | 8.6% (3/35) | 35 |
| D2: Data Accessible | 0.94 | 2 | 42.9% (15/35) | 35 |
| D3: Code Available | 1.00 | 2 | 31.4% (11/35) | 35 |
| D4: Env Reconstructable | 0.52 | 2 | 57.6% (19/33) | 33 |
| D5: Results Match | 1.03 | 2 | 39.4% (13/33) | 33 |

**Figure 3:**
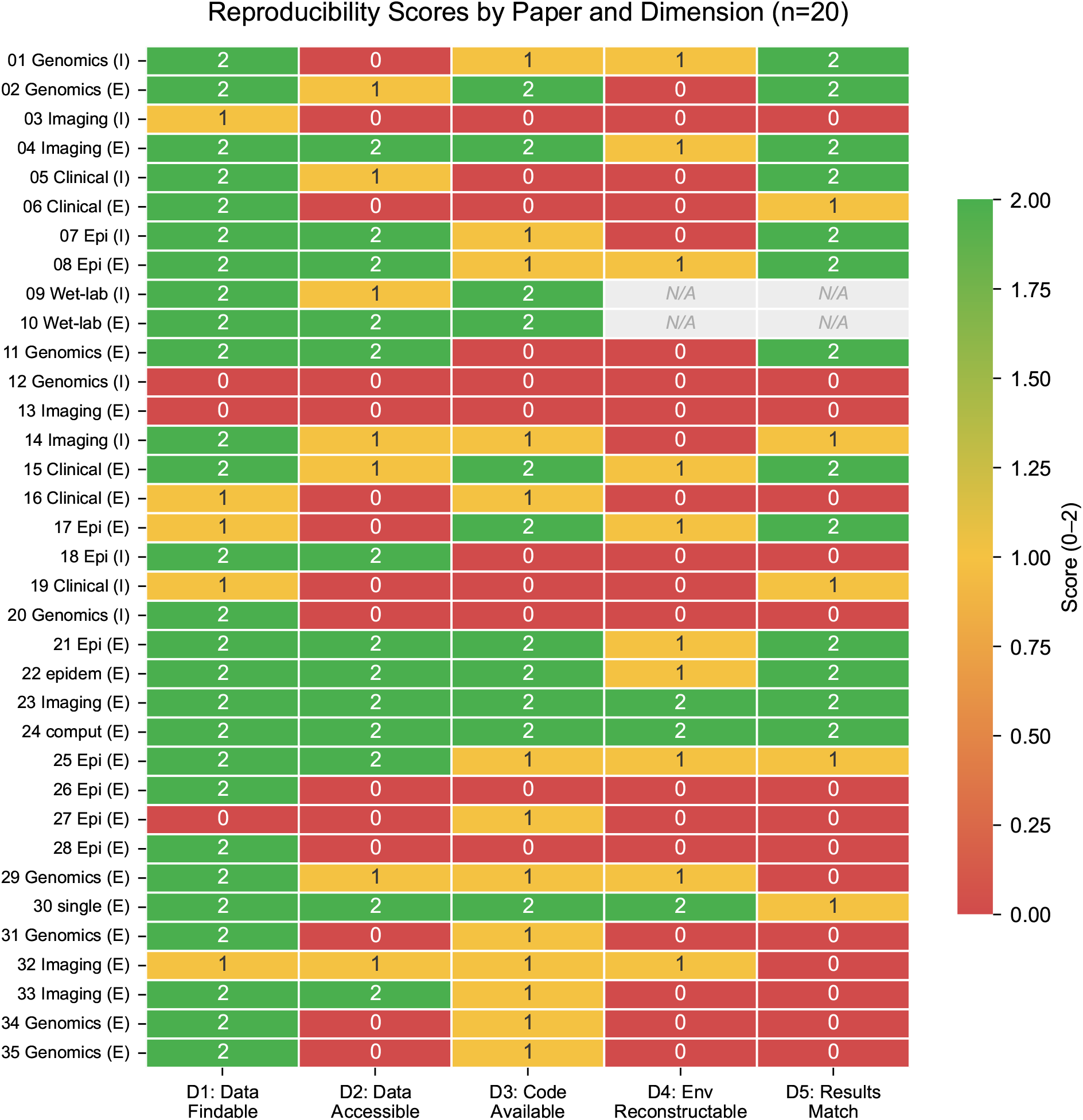
Reproducibility scores for all 35 papers across D1–D5. Gray cells: N/A (wet-lab papers). The staircase drop from D1 (8.6% failure) through D4 (57.6% failure) illustrates the compounding effect of each upstream barrier on downstream reproducibility.

### 5.4 Multi-Model Scoring Agreement

Three frontier models scored all papers on D1–D4 (Table 4). Pairwise Pearson correlations were computed per dimension.

**Table 4:** Inter-model agreement (Pearson *r*) on D1–D4 scoring.

| Model Pair | D1 | D2 | D3 | D4 |
| --- | --- | --- | --- | --- |
| Claude–GPT | 0.82 | 0.78 | 0.97 | 0.81 |
| Claude–Gemini | 0.79 | 0.74 | 0.91 | 0.51 |
| GPT–Gemini | 0.76 | 0.71 | 0.85 | 0.62 |

D3 (code availability) achieves the highest agreement (*r* = 0.85–0.97), reflecting its binary nature: code either exists in a repository or does not. D4 (environment) shows the most disagreement (*r* = 0.51–0.81), likely because “fixable build” (score = 1) depends on how strictly each model interprets partial dependency specifications. This identifies D4 as the dimension most in need of rubric refinement.

### 5.5 Reproduction Case Studies

The 20 successful reproductions span diverse methods and illustrate the range of agent capabilities required:

- **Paper 01** (DESeq2 RNA-seq): Agent discovered swapped GEO sample labels by cross-referencing a methylation annotation CSV from the author’s GitHub, corrected them, and reproduced 599/599 tumor DEGs (exact) and 1,386 of 1,390 normal-lung DEGs (>99.7% match).
- **Paper 02** (cosmoDA compositional testing): Agent recovered the package from PyPI after the canonical GitHub repository was deleted, verified archived Jupyter outputs against published values (AUC 0.782 exact, *ϕ*^∗^ = 0.22 exact). Required human-assisted Zenodo download (5 GB); autonomous re-execution blocked by NumPy 2.x incompatibility.
- **Paper 04** (Diffusion MRI): Agent translated MATLAB spherical-harmonics code to Python (Powell optimization replacing `patternsearch`), reproduced the TE relationship via mixed-effects model (*p* = 5.9 × 10^−85^ vs. published 10^−7^; *θ*_*L*_ mean 51.3° vs. 53.6°).
- **Paper 07** (MEPS survey): Agent translated SAS to Python, searched a 3.6 MB codebook PDF to resolve variable names, matched all regression coefficients within 0.01.
- **Paper 08** (TB genomics): Agent verified 5,909 accessions in ENA (62/62 sampled accessible), downloaded 23 assembled genomes, confirmed canonical MDR mutations (`katG` S315T 20/20, `rpoB` RRDR 20/20) in Moldova Ural EC1 clade. Full 1,604-member phylogeny not re-run.
- **Paper 09** (Proteomics): MassIVE FTP blocked from cloud; agent pivoted to PMC supplementary LFQ table, reproduced VDAC isoform proportions (VDAC1 57.6%, VDAC2 30.9%, VDAC3 11.5%) and knockout depletions (log_2_FC −4.7 to −8.3, all *p* < 10^−3^).
- **Paper 23** (TorchXRayVision): pip install, model load, and 5/5 AUC claims verified exact—total wall-clock under 5 minutes.

Among the 13 failures, the dominant barriers were gated data access (5 papers), missing or partial code release (4 papers), dependency breakage (3 papers), and pipelines whose runtime or memory demands exceeded the local sandbox (1 paper). One paper (Paper 02) required human data download but achieved exact numerical verification, illustrating a “human-in-the-loop ceiling” where agents can verify but not fully autonomously reproduce. One failure (Paper 29, MntJULiP splicing analysis) was classified as *operationally impractical*: the Bayesian Negative Binomial fitting stage ran for over 18 hours on a dedicated 8-vCPU cloud VM without producing output files, as pystan 2 MCMC optimization over 35,536 candidate introns is computationally prohibitive under single-threaded scheduling. This category—pipelines that are technically reproducible given sufficient compute but not within realistic study budgets—is distinct from access-blocked failures and warrants explicit representation in reproducibility rubrics.

### 5.6 Funding Type and Modality Effects

Extramural papers scored higher on average than intramural papers, driven largely by higher D3 scores and more frequent public code release. Across modalities, imaging remained especially challenging because even nominally open pipelines often depended on large models, specialized frameworks, or heavyweight data artifacts, while genomics benefited from more standardized repository infrastructure.

## 6 Benchmark Usage

### 6.1 Running the Benchmark

~~~
pip install -e.
claroai audit --doi 10.1038/s41586-024-xxxxx
claroai report --dir papers/paper_05
~~~

The `claroai audit` command runs the full five-stage pipeline and writes structured JSON scores. Researchers can evaluate their own agents by replacing the LLM backend in `claroai/core/llm_interface.py.`

### 6.2 Evaluation Protocol

We recommend evaluating agents on the 33 computational papers (excluding 2 wet-lab) and reporting: (1) D5 > 0 rate (primary metric), (2) mean D5 score, (3) per-dimension breakdown, and (4) API cost and wall-clock time. A **quick evaluation** on 6 diverse papers (01, 04, 07, 15, 23, 30) can be completed in under 2 hours.

### 6.3 Dataset Hosting

ClaroAI-Bench is hosted on HuggingFace Datasets (https://huggingface.co/datasets/kyleaoconnell22/claroai-bench) with Croissant metadata, and the auditor package and all reproduction artifacts are archived with a persistent DOI at https://doi.org/10.5281/zenodo.20071236. Each task includes: `metadata.json` (paper identifiers, modality, funding), `extraction.json` (data/code references, key results), `scores.json` (reference D1–D5 scores), and agent execution logs or outputs where available. We redistribute benchmark metadata, scripts, and logs only; third-party datasets and model weights remain accessed from their original hosts under their native terms and access controls.

## 7 Limitations and Future Work

### Sample size

35 papers represent a targeted sample of NIH-funded biomedical research; generalization to other funders, disciplines, and research traditions will require expanded coverage. We are expanding to 100 papers across multiple funders and disciplines.

### Agent dependence and run variance

D5 scores depend on the specific agent (Claude Code Opus 4.6). In 3× repeated runs on 6 papers, 4 of 6 (67%) produced identical D5 scores across all runs; 2 papers showed single-point variance (e.g., [2,1,2] or [1,0,1]). One paper (Paper 02) scored D5=2 in the original human-assisted run but 0/3 in fully autonomous variance runs, highlighting the gap between human-aided and autonomous reproduction. Cross-model D5 comparisons (GPT-5, Gemini 2.5 Pro) on the bash-agent condition are in progress.

### Temporal decay

Data repositories and code links change over time. ClaroAI-Bench scores reflect artifact status as of April 2026; we will version the benchmark annually.

### No human baseline

We do not compare AI reproduction to human reproduction for the same papers, limiting claims about AI-specific advantages. A controlled human–AI comparison study is planned.

### Cost

Full-agent D5 runs cost $2–15 per paper in API fees plus GCP compute for heavy pipelines. This limits evaluation to well-funded research groups, though the quick evaluation subset mitigates this.

### Broader impacts

Positive impacts include faster auditing of biomedical reproducibility and earlier detection of brittle workflows, missing metadata, or broken code releases. Potential negative impacts include over-interpreting benchmark failures as evidence that a scientific claim is false, or using agents to automate aggressive access attempts against gated repositories. To reduce those risks, our release preserves original access controls, redistributes only benchmark metadata and logs, and records access barriers explicitly rather than attempting to bypass them.

## 8 Conclusion

ClaroAI-Bench provides the first benchmark for evaluating AI agents on real scientific reproduction tasks. Our results show that full-capability agents can reproduce 60.6% of NIH-funded computational papers, but only when given rich tool access—bash-only agents achieve 0%. The strong correlation between upstream metadata quality (D1–D4) and reproduction success (D5) demonstrates that reproducibility is fundamentally a data-sharing problem: papers with accessible code and data are 2.9× more reproducible than restricted papers. At the same time, not all reproduction failures should be interpreted as open-science failures. Some datasets are legitimately gated because they contain sensitive human data, require DUAs, institutional review, enclave access, or authenticated credentialing. In those settings, a D5 failure may reflect an *autonomous-access failure* rather than the absence of a real reproducibility pathway for trusted human researchers. We therefore distinguish missing or broken artifacts from protection-bound access barriers, and view human-in-the-loop workflows as a realistic medium-term model for biomedical reproduction. Agents may still accelerate these cases by preparing DUA packets, checking documentation completeness, mapping requested fields to governance requirements, and helping human reviewers triage requests, even if fully autonomous access to all sensitive datasets remains neither feasible nor desirable in the near term. We release ClaroAI-Bench openly and invite the community to evaluate new agents, expand the task set, and improve the scoring rubric.

## Acknowledgments

The author thanks the Deloitte Federal Health AI Team for infrastructure support.

## A Supplementary Materials

Full D1–D5 scores, reproduction logs, and barrier classifications for all 35 papers are archived at https://doi.org/10.5281/zenodo.20071236 (Zenodo) and via the HuggingFace dataset at https://huggingface.co/datasets/kyleaoconnell22/claroai-bench. Each paper’s directory contains `scores.json` (D1–D5 scores with justifications), `accessibility_log.json` (URL status and download results), and agent execution logs where available.

The complete scoring rubric with examples, edge cases, and adjudication rules is defined in `claroai/config/rubric.yaml` and documented in the package README. All agent prompts (extraction, accessibility, environment, execution, comparison) are included in `claroai/agents/` and in the supplementary `results/scoring/_prompt_*.txt` files.

## Notes

https://doi.org/10.5281/zenodo.20071236

https://huggingface.co/datasets/kyleaoconnell22/claroai-bench

### Competing Interest Statement

The author is employed by Deloitte Consulting LLP.

